# A Near Complete Zonal Map of Mouse Olfactory Receptors

**DOI:** 10.1101/234187

**Authors:** Longzhi Tan, X. Sunney Xie

## Abstract

In the mouse olfactory system, spatially regulated expression of > 1,000 olfactory receptors (ORs) ― a phenomenon termed “zones” ― forms a topological map in the main olfactory epithelium (MOE). However, the zones of most ORs are currently unknown. By sequencing mRNA of 12 isolated MOE pieces, we mapped out zonal information for 1,033 OR genes with an estimated accuracy of 0.3 zones, covering 81% of all intact OR genes and 99.4% of total OR mRNA abundance. Zones tend to vary gradually along chromosomes. We further identified putative non-OR genes that may exhibit zonal expression.

## Main Text

The olfactory system relies on a topological map of olfactory receptor (OR) expression. The first stage of map formation occurs in the main olfactory epithelium (MOE), where different OR genes are expressed in different yet partially overlapping “zones”. The zones were first discovered in rodents (Ressler et al., 1993; Vassar et al., 1993), and later observed in fish (Weth et al., 1996), insects (Vosshall et al., 1999), and primates (Horowitz et al., 2014). Zonal expression is crucial for downstream projection to the main olfactory bulb (Mombaerts et al., 1996; Ressler et al., 1994; Vassar et al., 1994), and may contribute to the establishment of the “one-neuron-one-receptor” rule (Chess et al., 1994; Hanchate et al., 2015; Malnic et al., 1999; Tan et al., 2015). However, existing zonal information either had a limited resolution (Zhang et al., 2004) or was available only for a small subset of < 100 ORs (Miyamichi et al., 2005), and was limited for non-OR genes (Duggan et al., 2008; Gussing and Bohm, 2004; Ling et al., 2004; Miyawaki et al., 1996; Norlin et al., 2001; Oka et al., 2003; Tietjen et al., 2005; Tietjen et al., 2003; Vedin et al., 2009; Whitby-Logan et al., 2004; Yoshihara et al., 1997). In this work, we generated a near complete zonal map of 1,033 mouse ORs by mRNA sequencing of isolated MOE pieces, and identified novel non-OR genes that may exhibit zonal expression.

To investigate the spatial pattern of gene expression in the mouse MOE, we sequenced mRNA from 12 isolated MOE pieces with an average of 12.5 million single-end 100-bp reads per piece (min = 4.9 million, max = 21.0 million) (**Table S1–S3**). In mice, the expression zone of each OR assumes a complex shape that approximates a cylindrical shell, concentric to each other (Ressler et al., 1993). Here we follow a naming convention where zone 1 is the most dorsomedial, zone 5 (also known as zone 4b) is the most ventrolateral, and the zone of each OR is not necessarily an integer (Miyamichi et al., 2005) (**Figure 1A**). The geometry is further complicated by an unusual, medial zone of *Olfr459* (also known as OR-Z6 or MOR120-1, orthologous to human OR9A2) (Pyrski et al., 2001) (red region in **Figure 1A**). We tackle the problem of the complex zonal geometry and its invisibility to naked eyes by randomly isolating 12 very small MOE pieces, each of which would contain only one or a few nearby zones. In this way, the expression pattern of each OR across the 12 MOE pieces would reflect its zone. Indeed, normalized expression levels of the 1,033 OR genes that were sufficiently detected (defined as having an average of at least 0.5 transcripts per million (TPM)) showed a prominent pattern of differential expression across the 12 pieces, as visualized by t-distributed stochastic neighbor embedding (t-SNE) (Maaten and Hinton, 2008) and principal component analysis (PCA) (black points in **Figure 1B**). This pattern reflected zonal expression, because on the t-SNE and PCA plots a small subset of 78 ORs of known zones (Miyamichi et al., 2005; Pyrski et al., 2001) were distributed according to their zones (colored points in **Figure 1B**) (**Table S4**). For example, on the t-SNE plot, *Olfr459* of the unusual zone resided in a small, separate cluster at the top, while the other 77 ORs of zones 1 to 5 followed a continuous “trajectory” along the larger cluster.

**Figure 1.**
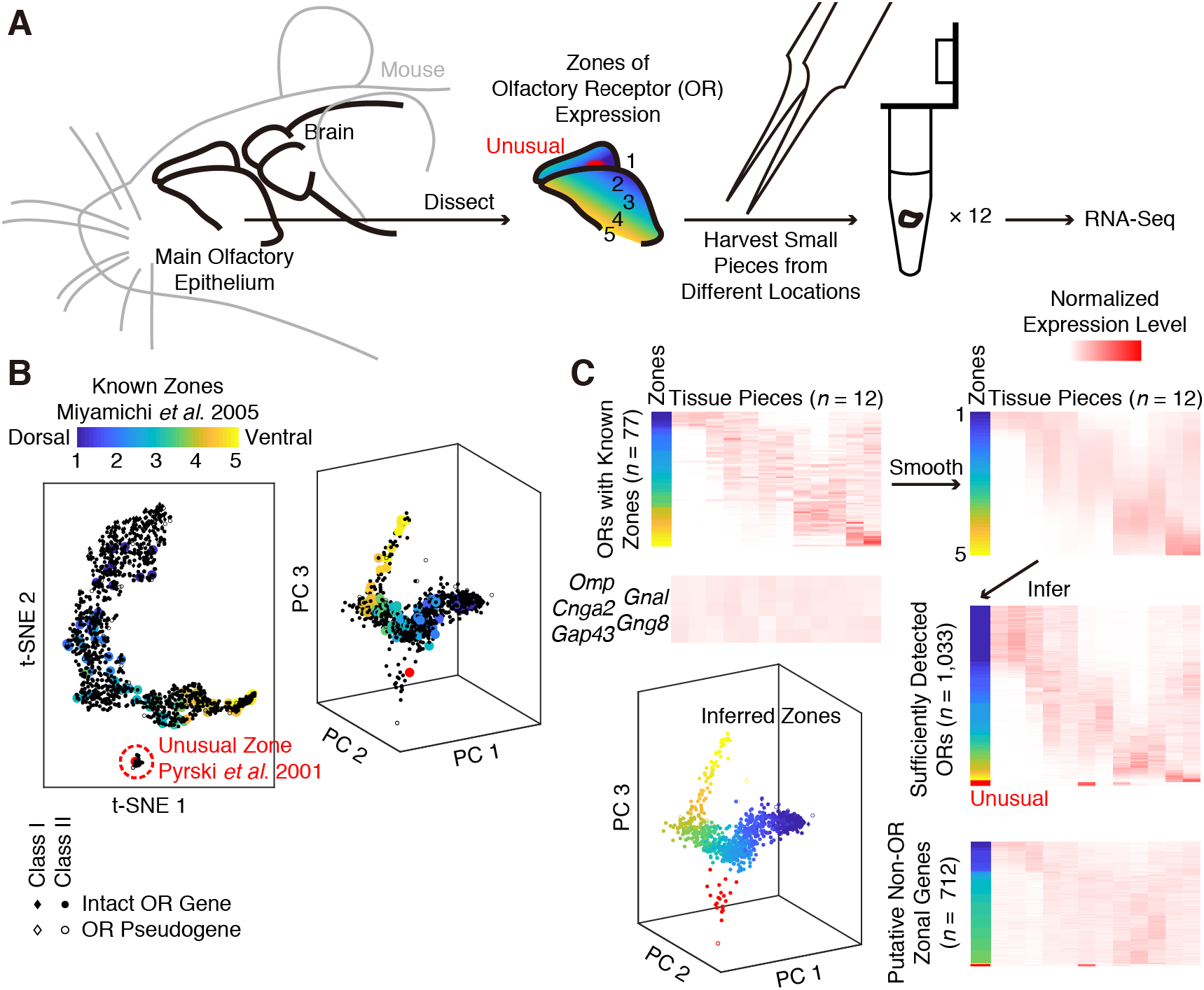
Spatial-transcriptomic mapping of the expression zones of 1,033 olfactory receptor (OR) genes and 712 putative non-OR genes. (**A**) We investigated the spatial pattern of gene expression in the mouse main olfactory epithelium (MOE) by sequencing mRNA from 12 randomly isolated MOE pieces. Each small piece would contain only one or a few nearby zones. (**B**) Normalized expression levels of the 1,033 OR genes (diamonds for Class I ORs (Zhang and Firestein, 2002) and dots for Class II) that were sufficiently detected showed a prominent pattern of zonal expression across the 12 pieces, as visualized by 78 ORs of known zones (colored points) (Miyamichi et al., 2005) via t-distributed stochastic neighbor embedding (t-SNE) and principal component analysis (PCA). We assigned the 22 ORs in the smaller t-SNE cluster (red dashed circle) to the unusual zone of *Olfr459* (red dot). (**C**) We made a standard curve from zone 1 to zone 5 by smoothing the normalized expression levels of the 77 known ORs (top heatmaps), and inferred zones of the 1,033 ORs by finding the closest point on the standard curve (middle right heatmap). Results of the inference was visualized on a PCA plot. Putative non-OR genes that may exhibit zonal expression were selected based on expression levels, uniformity across the MOE pieces, and distance to the standard curve or to *Olfr459* (bottom heatmap). In contrast, known marker genes (*Omp, Gnal, Cnga2* for mature neurons, and *Gng8, Gap43* for immature neurons) exhibited uniform expression (middle left heatmap) .

We inferred the expression zones of 992 intact OR genes (81% of all 1,228, based on the GENCODE annotation) and 41 OR pseudogenes (22% of all 189) using the observed expression patterns of 78 known ORs as standards. Among the 1,033 ORs that were sufficiently detected, we began by assigning the 22 ORs in the smaller t-SNE cluster (**Figure 1B**) to the unusual zone (Pyrski et al., 2001), showing for the first time that the unusual expression pattern of *Olfr459* is not an isolated case. For the remaining ORs, which would belong to the usual zones, we first made a 12-dimensional standard curve from zone 1 to zone 5 by smoothing the normalized expression levels of the 77 known ORs (Miyamichi et al., 2005) with a half window size of ± 0.5 zones (top heatmaps in **Figure 1C**). The zone of each OR was then inferred by finding the closest point (as measured by Euclidean distance) on the standard curve (middle heatmap and visualization on a PCA plot in **Figure 1C**) (**Table S5**). Uncertainties in zonal inference could be estimated by leave-one-out cross-validation on the 77 known ORs, yielding a root-mean-square error of 0.3 zones (min = 0.0, max = 1.1). Together, these ORs accounted for 99.4% of the total OR mRNA abundance (as measured by average TPMs) in the MOE.

We further identified 666 intact non-OR genes and 46 non-OR pseudogenes that may exhibit zonal expression. The large number of annotated genes in the mouse genome necessitated stringent criteria. We removed genes whose expression was either low (average TPM < 1) or uniform across the 12 pieces (coefficient of variation < 0.5). Among the remaining 3,228 non- OR genes, putative zonal genes were selected based on Euclidean distance to the aforementioned standard curve (699 candidate genes with distance ≤ 2) or to the unusual zone’s *Olfr459* (13 genes with distance ≤ 4) (bottom heatmap in **Figure 1C, Table S4**). This list included known zonal genes *Acsm4* (also known as O-MACS) (Oka et al., 2003), *Nqo1* (Gussing and Bohm, 2004), *Ncam2* (also known as OCAM) (Yoshihara et al., 1997), *Foxg1* (Duggan et al., 2008), *Eya2* (Tietjen et al., 2003), *Msx1, Nrp2* (Norlin et al., 2001), *Gstm5* (Whitby-Logan et al., 2004), and trace-amine-associated receptors (TAARs) (Liberles and Buck, 2006). Novel genes included transcription factors *(Six3, Yy2, Isl1, Prdm16, Tbx3/15, Bach2, Foxa1, Pitx1, Tcea3/l3/l5, Npas3/4, Dlx3, Twist1/2, E2f2/8, Zfp97/365/382/950)*, chemokines (*Cxcl10/14*, *Ccl5/8*), cytochromes (Ling et al., 2004) (*Cyp2a4/2b10/2c44/2e1/7b1*), and aldehyde dehydrogenases (Norlin et al., 2001) (*Aldh1a7/3a1/3b1*). The false discovery rate (FDR) could be estimated to be 51% for the usual zones by random permutation of MOE-piece labels. A more stringent list of 202 genes (distance ≤ 1.5 to the standard curve) was also created (**Table S5**), with an estimated FDR of 18%. These genes provide promising candidates for zonal regulation of OR expression and of other cellular characteristics, for example cilia lengths (Challis et al., 2015).

Distribution of zones along the mouse genome provides insights into the mechanism of zonal regulation. Within each OR gene cluster, inferred zones typically vary gradually along the chromosome, although drastic changes between adjacent ORs occasionally occur (**Figure 2A**). In the canonical zones, two ORs would differ by an average of 0.3, 0.8, 0.9, or 1.3 zones, respectively, when they are separated by 10 kb, 100 kb, 1 Mb, or two different chromosomes (**Figure 2B**). This hints to a model where the pattern of chromatin silencing morphs continuously from zone to zone, in each zone exposing only a subset of ORs ― namely, the ORs of that zone ― to some universal machinery of transcriptional activation. This model is consistent with a recent observation that removal of H3K9-mediated heterochromatic silencing obliterated zonal expression of ORs, among other changes in OR expression (Lyons et al., 2014). In addition, genomic locations of drastic zonal changes may harbor important regulatory sites for OR silencing. Although most non-OR candidates reside in separate locations (**Figure 2C**), several are embedded in OR clusters and share zones with nearby ORs, likely a byproduct of chromatin organization under zonal regulation.

**Figure 2.**
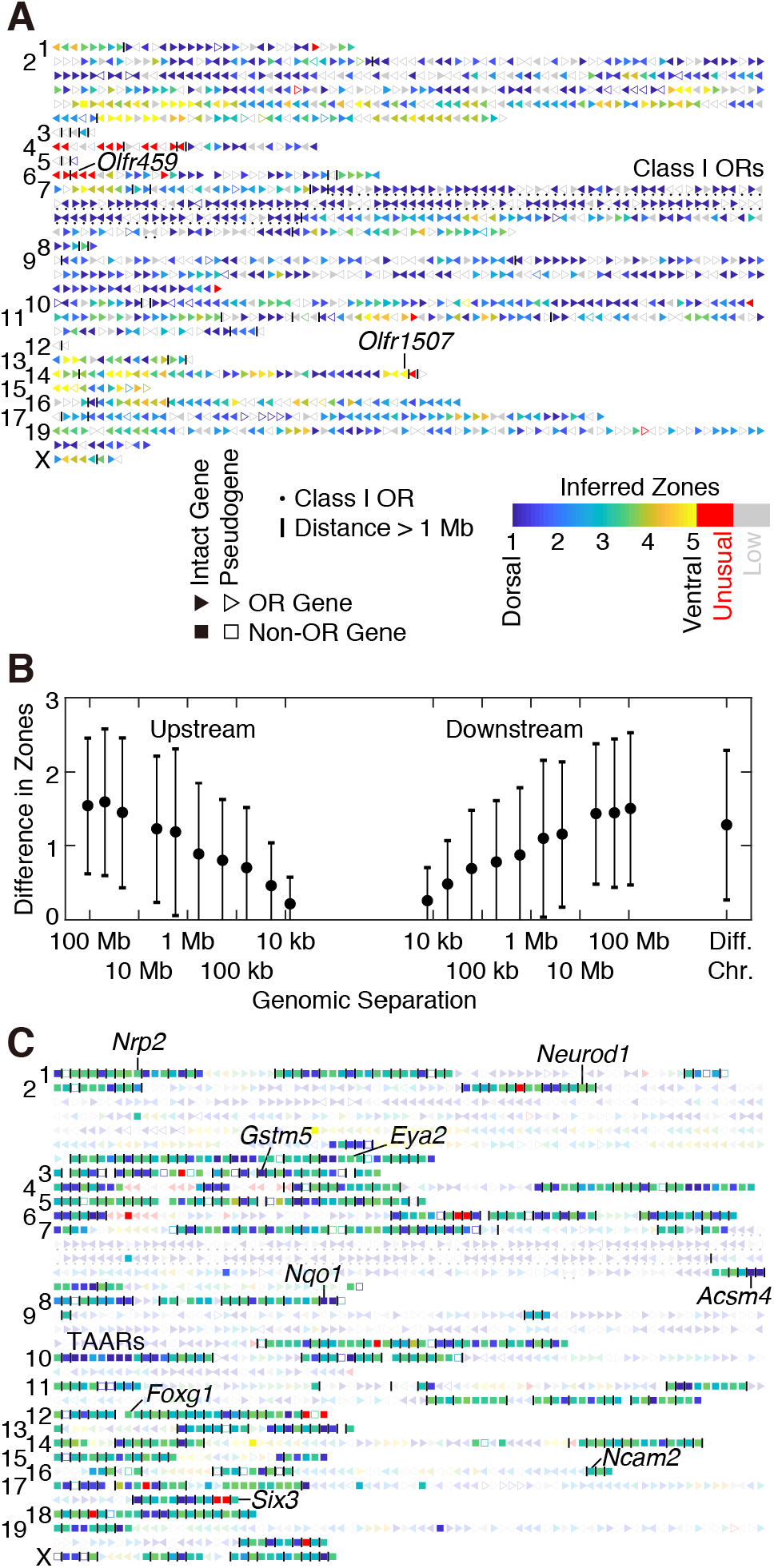
Distribution of zones across the mouse genome. (**A**) Zonal information for all 1,417 annotated ORs (triangles, with the pointy end denoting the direction of transcription). Typically, inferred zones vary gradually along the chromosome, although drastic changes between adjacent ORs occasionally occur. Numbers on the left denote chromosomes. Grey denotes insufficient detection because of low expression levels. OR clusters are separated by vertical lines. (**B**) Nearby ORs tend to have similar zones. All pairs of ORs from the canonical zones were grouped into bins according to their log10 genomic separation (in base pairs, upstream or downstream) with a bin size of 0.5. Bins with fewer than 10 OR pairs were excluded. Each black dot denotes the average difference in zones and the average log10 genomic separation in each bin. Error bar denotes standard deviation. (**C**) Zonal information for 712 non-OR candidates (squares), displayed together with ORs (triangles, faded for visual clarity).

Our results provide by far the most complete zonal map of mouse ORs, covering an order of magnitude more OR genes than state-of-the-art results obtained by traditional *in situ* hybridization (Miyamichi et al., 2005). Future work includes improving zonal resolution and inferring lowly expressed ORs by deeper sequencing of more MOE pieces, improving zonal inference by allowing variable kernel size and investigating expression patterns within zone 1, mapping out the second stage of the olfactory map by applying the same spatial-transcriptomic approach to the olfactory bulb, and comparing zonal expression between mouse and human.

## Materials and Methods

### Animals

All mouse experiments were performed in accordance with relevant guidelines and regulations. Animal protocols were approved by Harvard IACUC. An adult male mouse from the inbred strain C57BL/6NTac (Taconic) was used.

### Published Data

Known zonal information of 82 ORs was downloaded from Table S1 of (Miyamichi et al., 2005) and converted to modern gene names through MGI (**Table S4**), among which 77 were sufficiently detected in our experiment.

### RNA-Seq Experiments

Small pieces were isolated by forceps from a dissected MOE. Total RNA was extracted with an RNeasy Mini Kit (Qiagen) and a TissueLyser II (Qiagen) (25 Hz for 2 min, invert, and another 2 min, with 5 mm stainless steel beads), and quantified by an RNA 6000 Pico Kit on a Bioanalyzer 2100 (Agilent).

Libraries were prepared with a TruSeq RNA Sample Prep Kit v2 (Illumina) with poly(A) selection, quantified by a Qubit dsDNA HS Assay Kit (Thermo Fisher Scientific) and a High Sensitivity DNA Analysis Kit on a Bioanalyzer 2100 (Agilent), and sequenced on a HiSeq 2500 (Illumina). **Table S1** shows the number of reads in each library.

### Data Analysis

RNA-Seq reads were mapped to the mouse reference genome GRCm38.p5 (downloaded from the GENCODE M12 release) with tophat v2.0.11 (with default parameters) (Kim et al., 2013). Expression levels were quantified with cufflinks v2.2.1 (with parameters “-u--max-bundle-frags 100000000”) (Trapnell et al., 2010) based on the comprehensive gene annotation (ALL) from the above GENCODE release. Restricting quantification to reads that have the highest mapping quality (of 50) had no impact on the results. TPM values were calculated after removing genes that have a gene type beginning with “Mt_”, “miRNA”, “rRNA”, “scRNA”, “snRNA”, “snoRNA” “sRNA”, “scaRNA”, or “vaultRNA”. Translated protein sequences were downloaded from the above GENCODE release. Gene descriptions were downloaded from Ensembl BioMart (mouse genes GRCm38.p5). OR genes (1,417 genes) were defined as having “Olfr” in the gene name (1,419 entries, 4 pairs of which refer to the same genes: ENSMUSG00000109148 and ENSMUSG00000074985 are both *Olfr1452-ps1*, ENSMUSG00000109806 and ENSMUSG00000073919 are both *Olfr663*, ENSMUSG00000109020 and ENSMUSG00000061501 are both *Olfr197*, and ENSMUSG00000063732 and ENSMUSG00000110991 are both *Olfr908*) or having “olfactory receptor” in the gene description (an additional 2 genes, “Olrf445-ps1” and “OR4P4”). Pseudogenes (5,580 genes, 189 of which are ORs) were defined as having “pseudogene” in the gene type.

For zonal inference, expression levels (in TPMs) were normalized such that the average is 1 for each gene across the 12 MOE pieces. Two-dimensional t-SNE was performed with its MATLAB implementation on pairwise Euclidean distances between normalized expression values (the “tsne_d” function, with a default perplexity of 30). In the main text, the average difference in zones given a genomic separation was calculated from all OR pairs with log_10_ genomic separations within ± 0.1 of the desired value, regardless of directionality (upstream or downstream).

### Data Availability

Raw sequencing data were deposited at the National Center for Biotechnology Information with accession number SRP127539 at the following link: http://www.ncbi.nlm.nih.gov/sra/SRP127539

## Acknowledgments

The authors thank Stavros Lomvardas and Kevin Monahan (Columbia University), Chenghang Zong (Baylor College of Medicine), Catherine Dulac (Harvard University), Yang Shi (Boston Children’s Hospital), Stephen Liberles, Qian Li, Sean Megason, and Tim Mitchison (Harvard Medical School) for helpful discussion. This work was supported by an NIH Transformative Research Award (R01 EB010244) and an NIH Director’s Pioneer Award (DP1 CA186693) to X.S.X.. L.T. was supported by an HHMI International Student Research Fellowship.

## Author Contribution

L.T. and X.S.X. designed the experiments. L.T. carried out experiments and analyzed data.

## Conflict of Interest

The authors declare that they have no conflict of interest.

---

**Table S1.** Number of sequencing reads for the 12 MOE pieces.

**Table S2.** Expression levels in FPKMs of all 49,978 genes across the 12 MOE pieces, generated by combining Cufflinks outputs.

**Table S3.** Expression levels in TPMs of all 44,434 genes across the 12 MOE pieces, calculated from FPKMs after filtering out undesired genes (see Methods).

**Table S4.** Known zonal information for 82 ORs from (Miyamichi et al., 2005), with old gene names converted to modern ones.

**Table S5.** Inferred zones for all 1,417 ORs, corresponding to **Figure 2A**.

**Table S6.** Inferred zones for 712 non-OR genes that are putatively zonal, interleaved with all 1,417 ORs, corresponding to **Figure 2C**.

**Table S7.** Inferred zones for a more stringent set of 215 non-OR genes that are putatively zonal, interleaved with all 1,417 ORs.

## References

Challis, R.C., Tian, H., Wang, J., He, J., Jiang, J., Chen, X., Yin, W., Connelly, T., Ma, L., Yu, C.R., et al. (2015). An Olfactory Cilia Pattern in the Mammalian Nose Ensures High Sensitivity to Odors. Current biology: CB 25, 2503–2512.

Chess, A., Simon, I., Cedar, H., and Axel, R. (1994). Allelic Inactivation Regulates Olfactory Receptor Gene-Expression. Cell 78, 823–834.

Duggan, C.D., DeMaria, S., Baudhuin, A., Stafford, D., and Ngai, J. (2008). Foxg1 is required for development of the vertebrate olfactory system. The Journal of neuroscience: the official journal of the Society for Neuroscience 28, 5229–5239.

Gussing, F., and Bohm, S. (2004). NQO1 activity in the main and the accessory olfactory systems correlates with the zonal topography of projection maps. Eur J Neurosci 19, 2511–2518

Hanchate, N.K., Kondoh, K., Lu, Z., Kuang, D., Ye, X., Qiu, X., Pachter, L., Trapnell, C., and Buck, L.B. (2015). Single-cell transcriptomics reveals receptor transformations during olfactory neurogenesis. Science 350, 1251–1255.

Horowitz, L.F., Saraiva, L.R., Kuang, D., Yoon, K.H., and Buck, L.B. (2014). Olfactory receptor patterning in a higher primate. The Journal of neuroscience: the official journal of the Society for Neuroscience 34, 12241–12252.

Kim, D., Pertea, G., Trapnell, C., Pimentel, H., Kelley, R., and Salzberg, S.L. (2013). TopHat2: accurate alignment of transcriptomes in the presence of insertions, deletions and gene fusions. Genome biology 14, R36.

Liberles, S.D., and Buck, L.B. (2006). A second class of chemosensory receptors in the olfactory epithelium. Nature 442, 645–650.

Ling, G., Gu, J., Genter, M.B., Zhuo, X., and Ding, X. (2004). Regulation of cytochrome P450 gene expression in the olfactory mucosa. Chem Biol Interact 147, 247–258.

Lyons, D.B., Magklara, A., Goh, T., Sampath, S.C., Schaefer, A., Schotta, G., and Lomvardas, S. (2014). Heterochromatin-mediated gene silencing facilitates the diversification of olfactory neurons. Cell reports 9, 884–892.

Maaten, L.v.d., and Hinton, G. (2008). Visualizing data using t-SNE. Journal of Machine Learning Research 9, 2579–2605.

Malnic, B., Hirono, J., Sato, T., and Buck, L.B. (1999). Combinatorial receptor codes for odors. Cell 96, 713–723.

Miyamichi, K., Serizawa, S., Kimura, H.M., and Sakano, H. (2005). Continuous and overlapping expression domains of odorant receptor genes in the olfactory epithelium determine the dorsal/ventral positioning of glomeruli in the olfactory bulb. The Journal of neuroscience: the official journal of the Society for Neuroscience 25, 3586–3592.

Miyawaki, A., Homma, H., Tamura, H., Matsui, M., and Mikoshiba, K. (1996). Zonal distribution of sulfotransferase for phenol in olfactory sustentacular cells. EMBO J 15, 2050–2055.

Mombaerts, P., Wang, F., Dulac, C., Chao, S.K., Nemes, A., Mendelsohn, M., Edmondson, J., and Axel, R. (1996). Visualizing an olfactory sensory map. Cell 87, 675–686.

Norlin, E.M., Alenius, M., Gussing, F., Hagglund, M., Vedin, V., and Bohm, S. (2001). Evidence for gradients of gene expression correlating with zonal topography of the olfactory sensory map. Molecular and cellular neurosciences 18, 283–295.

Oka, Y., Kobayakawa, K., Nishizumi, H., Miyamichi, K., Hirose, S., Tsuboi, A., and Sakano, H. (2003). O-MACS, a novel member of the medium-chain acyl-CoA synthetase family, specifically expressed in the olfactory epithelium in a zone-specific manner. Eur J Biochem 270, 1995–2004.

Pyrski, M., Xu, Z., Walters, E., Gilbert, D.J., Jenkins, N.A., Copeland, N.G., and Margolis, F.L. (2001). The OMP-lacZ transgene mimics the unusual expression pattern of OR-Z6, a new odorant receptor gene on mouse chromosome 6: implication for locus-dependent gene expression. The Journal of neuroscience: the official journal of the Society for Neuroscience 21, 4637–4648.

Ressler, K.J., Sullivan, S.L., and Buck, L.B. (1993).A zonal organization of odorant receptor gene expression in the olfactory epithelium. Cell 73, 597–609.

Ressler, K.J., Sullivan, S.L., and Buck, L.B. (1994). Information coding in the olfactory system: evidence for a stereotyped and highly organized epitope map in the olfactory bulb. Cell 79, 1245–1255.

Tan, L., Li, Q., and Xie, X.S. (2015). Olfactory sensory neurons transiently express multiple olfactory receptors during development. Mol Syst Biol 11, 844.

Tietjen, I., Rihel, J., and Dulac, C.G. (2005). Single-cell transcriptional profiles and spatial patterning of the mammalian olfactory epithelium. The International journal of developmental biology 49, 201–207.

Tietjen, I., Rihel, J.M., Cao, Y., Koentges, G., Zakhary, L., and Dulac, C. (2003). Single-cell transcriptional analysis of neuronal progenitors. Neuron 38, 161–175.

Trapnell, C., Williams, B.A., Pertea, G., Mortazavi, A., Kwan, G., van Baren, M.J., Salzberg, S.L., Wold, B.J., and Pachter, L. (2010). Transcript assembly and quantification by RNA-Seq reveals unannotated transcripts and isoform switching during cell differentiation. Nat Biotechnol 28, 511–515.

Vassar, R., Chao, S.K., Sitcheran, R., Nunez, J.M., Vosshall, L.B., and Axel, R. (1994). Topographic organization of sensory projections to the olfactory bulb. Cell 79, 981–991.

Vassar, R., Ngai, J., and Axel, R. (1993). Spatial segregation of odorant receptor expression in the mammalian olfactory epithelium. Cell 74, 309–318.

Vedin, V., Molander, M., Bohm, S., and Berghard, A. (2009). Regional differences in olfactory epithelial homeostasis in the adult mouse. The Journal of comparative neurology 513, 375–384.

Vosshall, L.B., Amrein, H., Morozov, P.S., Rzhetsky, A., and Axel, R. (1999). A spatial map of olfactory receptor expression in the Drosophila antenna. Cell 96, 725–736.

Weth, F., Nadler, W., and Korsching, S. (1996). Nested expression domains for odorant receptors in zebrafish olfactory epithelium. Proc Natl Acad Sci U S A 93, 13321–13326.

Whitby-Logan, G.K., Weech, M., and Walters, E. (2004). Zonal expression and activity of glutathione S-transferase enzymes in the mouse olfactory mucosa. Brain Res 995, 151–157.

Yoshihara, Y., Kawasaki, M., Tamada, A., Fujita, H., Hayashi, H., Kagamiyama, H., and Mori, K. (1997). OCAM: A new member of the neural cell adhesion molecule family related to zone-to-zone projection of olfactory and vomeronasal axons. The Journal of neuroscience: the official journal of the Society for Neuroscience 17, 5830–5842.

Zhang, X., and Firestein, S. (2002). The olfactory receptor gene superfamily of the mouse. Nat Neurosci 5, 124–133.

Zhang, X., Rogers, M., Tian, H., Zhang, X., Zou, D.J., Liu, J., Ma, M., Shepherd, G.M., and Firestein, S.J. (2004). High-throughput microarray detection of olfactory receptor gene expression in the mouse. Proc Natl Acad Sci U S A 101, 14168–14173.

